# Phylogenetic study of thermophilic genera *Anoxybacillus, Geobacillus, Parageobacillus,* and proposal of a new classification *Quasigeobacillus* gen. nov

**DOI:** 10.1101/2021.06.18.449068

**Authors:** Berenice Talamantes-Becerra, Jason Carling, Jochen Blom, Arthur Georges

## Abstract

A phylogenetic study of *Anoxybacillus, Geobacillus* and *Parageobacillus* was performed using publicly available whole genome sequences. A total of 113 genomes were selected for phylogenomic metrics including calculation of Average Nucleotide Identity (ANI) and Average Amino acid Identity (AAI), and a maximum likelihood tree was built from alignment of a set of 662 orthologous core genes. The combined results from the core gene tree and ANI and AAI dendrograms show that the genomes split into two main clades, clade I containing all *Geobacillus*, all *Parageobacillus* and some species of *Anoxybacillus*, and clade II, containing the majority of *Anoxybacillus* species. Clade I is further partitioned into three clades, consisting separately of *Geobacillus, Parageobacillus*, and a third clade which we suggest should be elevated to a new genus *Quasigeobacillus* gen. nov. Two species of *Anoxybacillus* showed inconsistent positioning among the trees produced by differing methods and could not be clearly resolved into any of the three existing genera or the new genus. This research shows the importance of considering closely related genera together when studying phylogeny or assigning genomic affinities.

## Introduction

Next-generation sequencing technologies produce massive amounts of information that allows researchers to discriminate among data to establish thresholds for bacterial taxonomical scales [1]. In recent decades, the use of DNA-DNA hybridization was considered as the “gold standard” for bacterial taxonomy. Alternatively, 16S rRNA gene sequencing has provided a different approach; however, its low-resolution results in high sequence similarities amongst bacterial genera and species, limiting its’ usefulness for classification [2]. As an alternative to the use of 16S rRNA gene sequencing for assessing bacterial populations, the average nucleotide identity (ANI) values provide a higher resolution to detect relatedness among bacterial genomes. It is calculated by comparing two closely related genome sequences and estimating the percentage identity of aligned nucleotides using the BLAST search program [3]. If comparisons are done between distant genomes, it is recommended to use the average amino acid identity (AAI), as amino acid similarity is better conserved in homologous protein sequences than DNA sequence similarity [4, 5]. The use of bioinformatic tools to calculate ANI by itself is not sufficient, as alignment of a set of orthologous core genes may provide a better view of the vertical phylogenetic relationship. Taxonomic affiliation of new genomes should be verified using average nucleotide identity and multilocus phylogenetic analysis [6].

The genera *Anoxybacillus, Geobacillus* and *Parageobacillus* belong to the family *Bacillaceae*. They are moderate thermophiles with the ability to form spores, allowing them to survive in a dormant state for years, depending on the environmental conditions of the host soil such as pH, water availability, organic matter or calcium. *Anoxybacillus* species have been isolated mainly from geothermal sources; however, they are also found as a food contaminant or in compost. *Geobacillus* species, have been isolated from hot springs, oilfields, spoiled canned food, desert sand, composts, water, ocean sediments, sugar beet juice and mud [7]. Similarly, *Parageobacillus* is a thermophilic facultative anaerobe, which has been isolated from geothermal springs environments and compost [8].

*Anoxybacillus* is a Gram-positive, spore-forming rods whose main fatty acid present in the membrane of is iso-C15:0, and has a reported range in genomic GC content 42–57 mol% [9]. *Anoxybacillus* was proposed in the year 2000 as a new genus within the family *Bacillaceae*, based on a strain isolated from animal manure. The anaerobic growth characteristics observed in this strain were reflected in the chosen generic name *Anoxybacillus*. The new strain, which was given the full name *Anoxybacillus pushchinensis*, was found to cluster together with the previously described *Bacillus flavothermus*, with both of these species being distinct from other *Bacillus* species. *Bacillus flavothermus* was then reclassified as *Anoxybacillus flavithermus* giving the new genus two species [10]. Over the subsequent years, numerous species and strains have been added to the genus *Anoxybacillus*, many of which have been found to be either aerobes or facultatively anaerobic [11, 12]. There are twenty-four species and subspecies belonging to genus *Anoxybacillus* validly described to date. The following thirteen have genome assemblies available: *A. amylolyticus* [13], *A. ayderensis* [14], *A. flavithermus* [10, 15], *A. flavithermus* subsp. *Yunnanensis* [16], *A. geothermalis* [17], *A. gonensis* [18], *A. kamchatkensis* [19], *A. mongoliensis* [20], *A. pushchinoensis* [10], *A. suryakundensis* [21], *A. tepidamans* [22], *A. thermarum* [23], *A. vitaminiphilus* [24]. The remaining eleven do not have genome assemblies available: *A. bogrovensis* [25], *A. caldiproteolyticus* [26, 27], *A. contaminans* [28], *A. eryuanensis* [29], *A. kamchatkensis* subsp. *Asaccharedens* [30], *A. kaynarcensis* [31], *A. kestanbolensis* [14], *A. rupiensis* [32], *A. salavatliensis* [33], *A. tengchongensis* [29] and *A. voinovskiensis* [34].

*Geobacillus* is a Gram-positive spore-forming thermophile, whose dominant cellular membrane fatty acids are iso-C15:0, iso-C16:0 and iso-C17:0, with a reported genomic GC content ranging between 48.2–58 mol% [9]. *Geobacillus* was also originally classified within the genus *Bacillus*. A taxonomic study of thermophilic *bacilli* in 2001 resulted in the splitting of a group from the genus *Bacillus* into the newly described genus *Geobacillus*, containing eight species, consisting of the two newly described species *Geobacillus subterraneus* and *Geobacillus uzenensis*, as well as the members previously belonging to *Bacillus* group 5: *G. stearothermophilus, G. thermoleovorans, G. thermocatenulatus, G. kaustophilus, G. thermoglucosidasius* and *G. thermodenitrificans* [35]. The species *G. thermoglucosidasius* was subsequently removed from the genus [36]. *Geobacillus* currently contains sixteen validly described species all of which have genome assemblies available. The remaining *Geobacillus* species described with a valid bacteriological code are: *G. icigianus* [37], *G. jurassicus* [38], *G. lituanicus* [39], *G. thermantarcticus* [27], *G. vulcani* [40], *G. genomospecies* 3 [36, 41], and *G. proteiniphilus* [42]. A genome sequence is available in Genbank with a species name of *G. zalihae* [43]; however, the name has not been validly published [9].

In 2016, the new genus *Parageobacillus* was defined by reassigning a clade from *Geobacillus*, integrating five species into this genus [36]: *P. caldoxylosilyticus* [44], *P. thermoglucosidasius* [27, 35, 45], *P. thermantarcticus* [27, 46], *P. toebii* [27, 47], *P. genomospecies 1* [36]. The taxonomic revision was based on placement of the clade within a core gene phylogenetic tree of *Geobacillus* species, GC content differences and ANI [36]. A partial phylogenetic analysis of *G. galactosidasius* by Ramaloko et al. [48], came to conclusion that *G. galactosidasius* should properly belong to *Parageobacillus*. Najar and Thakur [49] proposed the formal reclassification of *G. galactosidasius* to *P. galactosidasius* and *G. yumthangensis* to *P. yumthangensis*. The data from the present study agrees with this conclusion and we are following this reclassification for the taxonomy used in this study.

Here we report a phylogenetic analysis and comparative genomic analysis of the closely related thermophilic genera *Anoxybacillus, Geobacillus* and *Parageobacillus*, as well as determining the preliminary placement of 7 newly isolated strains of these genera [50]. The study was performed using 113 genome assemblies of *Anoxybacillus, Geobacillus* and *Parageobacillus* available on RefSeq database [51].

## Materials and methods

Methods followed in this paper were based on those used for reassessment of the genus *Geobacillus* by Aliyu et al. [36]. Specifically, a set of core genes was identified from the group of bacterial genomes under study, followed by construction of a maximum likelihood tree. The same set of genomes was used to calculate ANI and AAI scores, followed by construction of UPGMA dendrograms. GC content and genome size for all genomes were calculated. The topology of the trees, GC contents and ANI values were used for taxonomic delimitation [36].

### *Geobacillus* and *Anoxybacillus* genomes

Complete genome assemblies from the RefSeq database [51] were downloaded for thirty-two *Anoxybacillus* strains and seventy-three *Geobacillus* strains. The genus *Parageobacillus* was represented in the analysis by the genome assemblies of the seven validly described *Parageobacillus* species, with one representative assembly per species. Additionally, whole genome sequences of two *Anoxybacillus*, four *Geobacillus* and one *Parageobacillus* strains obtained from a study of thermophiles in Australia [50] were included. This group contained 31 chromosome level assemblies, 33 scaffold genomes and 49 genomes up to the contig level. Anomalous genome assemblies listed with suppressed RefSeq status, including those identified as chimeric, contaminated, or frameshifted were excluded from the study.

### Phylogenetic analysis

The *Anoxybacillus, Geobacillus, Parageobacillus* core genome was identified using EDGAR 2.0 [52]. The genome assembly of *Geobacillus thermodenitrificans* NG80-2 was used as a starting reference, from which all CDS were identified and iteratively compared against the set of selected *Anoxybacillus, Geobacillus* and *Parageobacillus* genome assemblies to identify orthologous genes via BLAST. This process results in the identification of a core genome consisting of the full set of CDS from the starting reference assembly, for which orthologs can be identified amongst all other assemblies [53]. The genome assembly of *Bacillus subtilis spizizenii* TU-B-10T [54] was included in the core gene analysis as an outgroup. The assemblies included are listed in **Supplementary Table S1**. During the core gene discovery process, five assemblies with relatively low numbers of CDS were excluded after they were found to significantly reduce the number of core genes recognised, and one was removed as it was an exact duplicate of another *Anoxybacillus Suryakundensis* strain. The assemblies removed are listed in **Supplementary Table S2**. The obtained CDS were concatenated and aligned using MUSCLE [55]. The alignments were used to build a maximum-likelihood tree with the software package FastTree [56, 57].

### Phylogenomic metric calculations

ANI and AAI values were calculated between all pairs of genomes to build ANI and AAI similarity matrices. Default parameters were selected to calculate two-way ANI using the ani.rb script; similarly, two-way AAI was calculated using the aai.rb script. Draft genomes were concatenated with 100 N bases introduced between contigs. Both Ruby scripts are publicly available in the enveomics package [5]. Similarity values obtained by ANI and AAI were arranged into similarity matrices and used as input in the DendroUPGMA [58] web server by selecting the unweighted pair group with arithmetic mean clustering method (UPGMA) to obtain dendrograms. Trees were plotted with the web-based tool Interactive Tree Of Life [59].

### Identifying genes unique to clade I-c

Genes unique to clade I-c were identified using BLASTn [60] alignment. To find the genes unique to clade I-C, CDS of clade I-c were used for BLASTn alignment against all the assemblies in the NCBI RefSeq database [51] and NCBI nt database. The relaxed BLASTn filtering criteria used were percentage identity 30%, bitscore 10 and percentage overlap of 1%. The CDS used for alignment were from the assembly *Anoxybacillus tepidamans* PS2, a member of clade I-c. Those CDS which were found to be present in all members of clade I-c and no members of clade I-a, clade I-b and clade II were considered as unique to clade I-c.

## Results

A total of thirty-two *Anoxybacillus*, seventy-three *Geobacillus*, eight *Parageobacillus* and one *Bacillus* genomes (**Supplementary Table S1**) produced a core gene set of 662 orthologous genes per genome, which was used to produce a maximum likelihood phylogenetic tree with 114 genomes.

The core gene maximum likelihood tree splits into three major groups consisting of the outgroup, clade I and clade II (**Figure 1**). Clade I consists of eighty-seven genomes, from *Geobacillus* and *Parageobacillus* plus a subset of *Anoxybacillus* genomes. Clade II contains twenty-four of the remainder *Anoxybacillus* genomes. Within clade I, the first branch separates *A. vitaminiphilus* from all other assemblies. The next branch splits the remaining *Anoxybacillus* with the exception of *A. flavithermus* B4168 from all *Geobacillus* and *Parageobacillus*. Clade I has been divided into *A. vitaminiphilus* plus three subclades, I-a, I-b and I-c.

**Figure 1.**
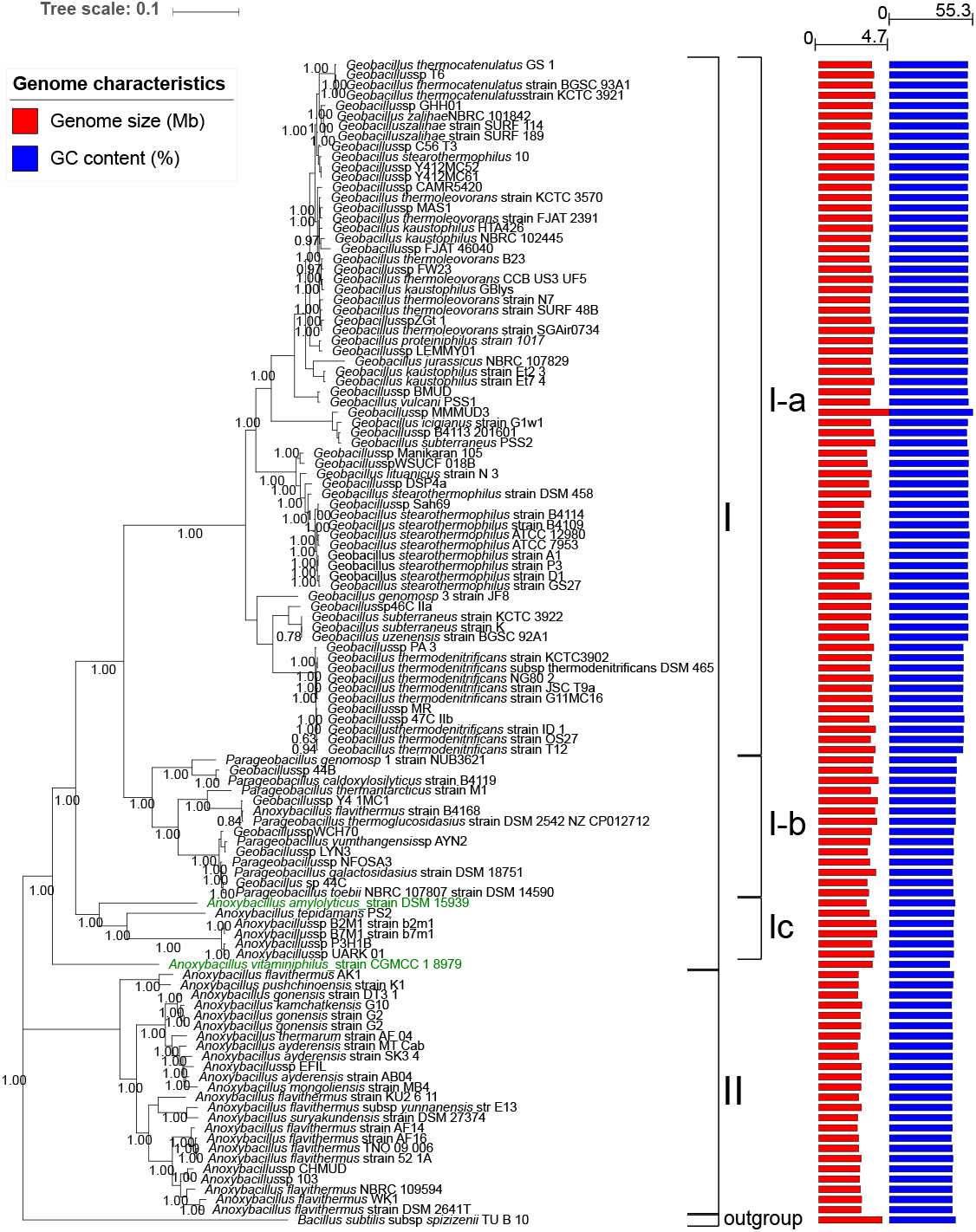

Clade I-a includes sixty-eight *Geobacillus* genomes from the species, *G. genomosp 3, G. icigianus, G. jurassicus, G. kaustophilus, G. lituanicus, G. stearothermophilus, G. subterraneus, G. thermocatenulatus, G. thermodenitrificans, G. thermoleovorans, G. uzenensis, G. vulcani, G. zalihae, G. proteiniphilus* and twenty-two unnamed *Geobacillus* species. Some species are represented by multiple genome assemblies.

*C*lade I-b contains fourteen genomes, from which seven representative species belong to *Parageobacillus* genomes including the species *P. genomosp 1, P. caldoxylosilyticus, P. thermantarcticus, P. thermoglucosidasius, P. toebii, P. galactosidasius, P. yumthangensis*. Five are unnamed *Geobacillus* and one unnamed *Parageobacillus* species. Additionally, the assembly named *A. flavithermus* B4168 is located within this clade.

Clade I-c has five genomes, from which one has been defined as *A. tepidamans*, and four are *unnamed Anoxybacillus* species.Clade II consists of twenty-four *Anoxybacillus* genomes, including the species *A. flavithermus, A. pushchinoensis, A. gonensis, A. kamchatkensis, A. thermarum, A. ayderensis,A. mongoliensis, A. suryakundensis and three unnamed Anoxybacillus* species. Some species are represented by multiple genome assemblies.

The two-way ANI values calculated between all genome pairs were used to build a UPGMA [58] dendrogram with 114 genomes (**Figure 2**). The clades from the maximum likelihood core gene tree are shown alongside for comparison. The first branch splits the outgroup from all other samples. The positions of *A. amylolyticus* and *A. vitaminiphilus* differ from the placement from the core gene maximum likelihood tree. The second branch splits *A. amylolyticus* from all other *Anoxybacillus, Geobacillus* and *Parageobacillus*, while *A. vitaminiphilus* is present within the clade of *Parageobacillus* (clade I-b). The ANI values calculated are shown in a similarity matrix in **Supplementary Table S3**.

**Figure 2.**
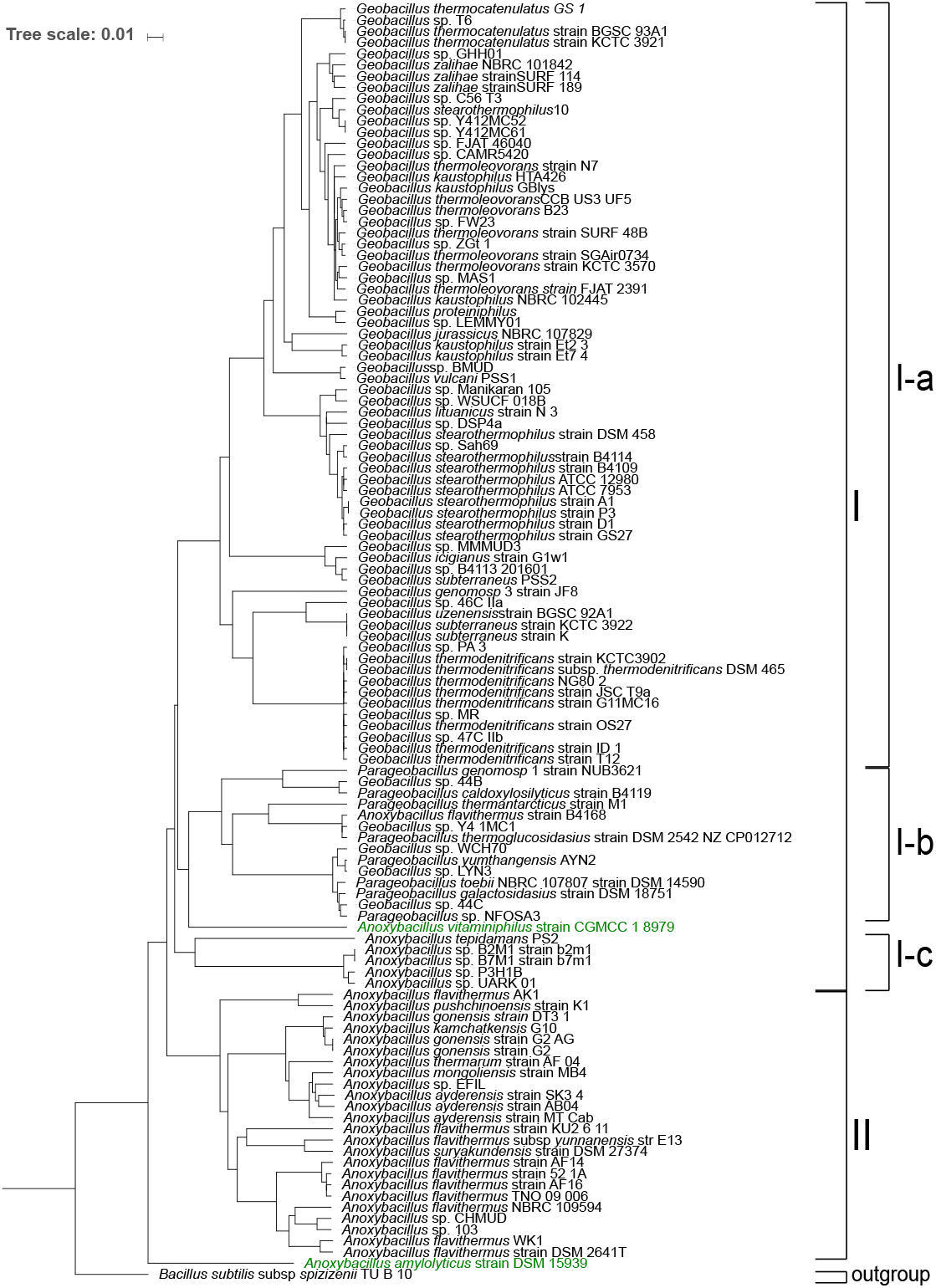

The two-way average AAI values calculated between all genome pairs were used to build a UPGMA [58] dendrogram with 114 genomes (**Figure 3**). The clades from the maximum likelihood core gene tree are shown alongside for comparison. Positioning is broadly similar to the trees in **Figure 1** and **Figure 2**. The primary distinction is the position of *A. amylolyticus*, and *A. vitaminiphilus* which are located in clade I-c within the subclade containing the other *Anoxybacillus* species. The AAI values calculated are shown in a similarity matrix in **Supplementary Table S4**.

**Figure 3.**
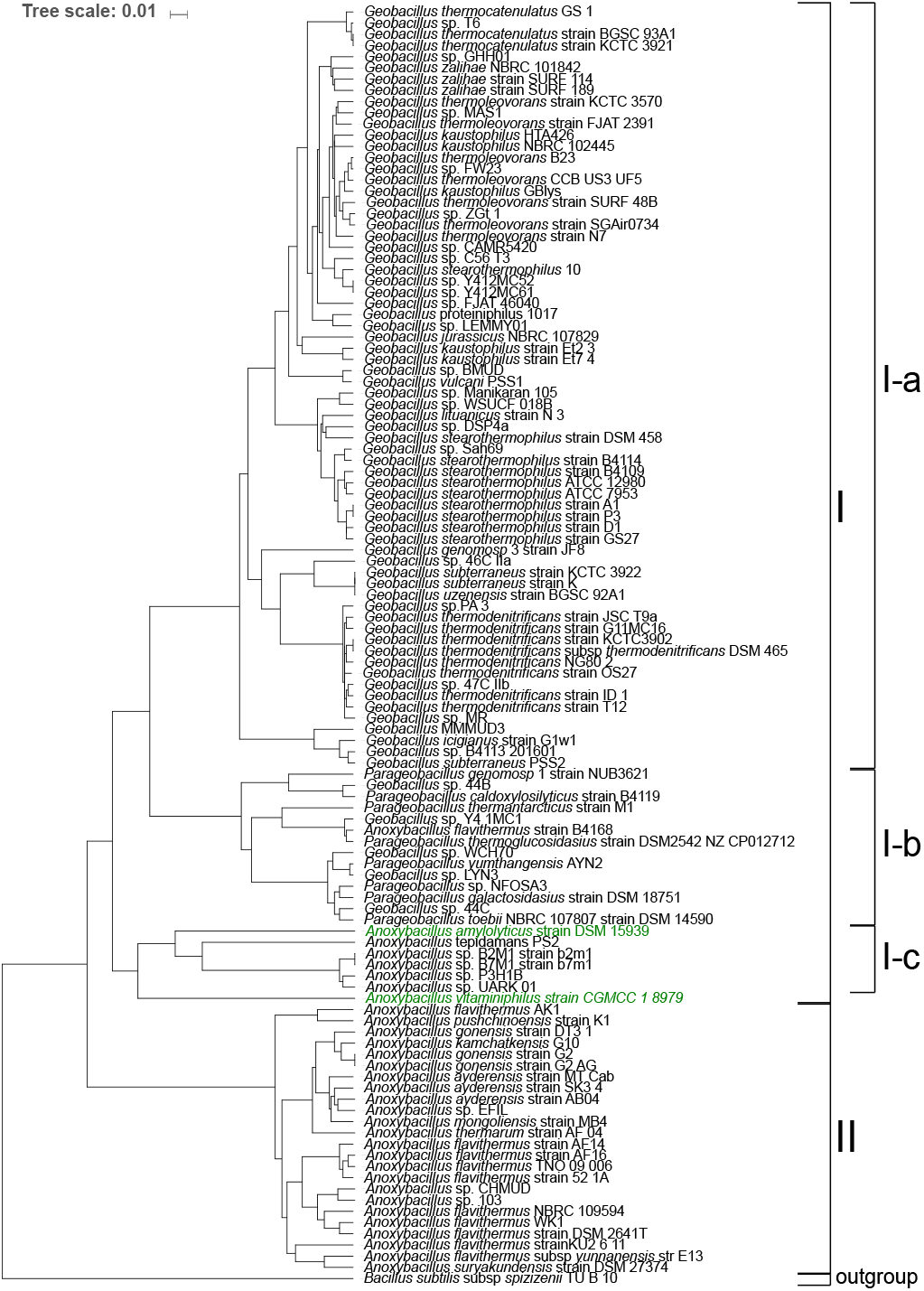

Genome size and GC content for each of the genomes is listed in **Supplementary Table S1**. A graphical representation of these values is shown alongside the core gene maximum likelihood tree in **Figure 1**.

The genome size of clade I-a contained mostly *Geobacillus* genomes ranges from 2.63 to 4.69 Mb and the GC content ranged from 48.79 to 55.27%.

Clade I-b, including *Parageobacillus* genomes, had genome sizes ranging from 3.22 to 3.95 Mb and GC content from 41.60 to 44.60%.

The clade I-c which includes a subset of *Anoxybacillus* genomes, showed genome sizes ranging between 3.36 to 3.87 Mb, and GC content ranging from 42.50 to 43.00%.

Clade II which includes the majority of *Anoxybacillus* genomes had a genome size range from 2.56 to 2.86 Mb and a GC content ranging from 41.10 to 42.70%.

### Affinities of seven newly sequenced genomes

The placement of the seven genomes derived from the study of thermophilic bacteria in Australia [50] was determined on the basis of their relative position within the core gene maximum likelihood tree and the ANI and AAI values. These genomes had genome sizes ranging between 2.73 to 4.69 Mb, a minimum GC content of 41.81% and a maximum of 55.27%. ANI and AAI values to their closest reference genomes, for the samples *Geobacillus* sp. BMUD, *Geobacillus* sp. MR, and *Parageobacillus* sp. NFOSA3 are above 99.17 and 98.39 respectively. Genomes of *Anoxybacillus* sp. EFIL, *Geobacillus* sp. MMMUD3 and *Geobacillus* sp. DSP4a have ANI of 97.99, 97.14 and 97.64 respectively. The genome of *Anoxybacillus* sp. CHMUD had the most distant ANI value with 95.03 and AAI of 95.81. Results are shown in **Table 3**.

**Table 1.** Phylogenomic metrics comparing the average nucleotide identity mean values and average amino acid identity mean values of candidate new genus assigned into clade I-c against the members of the existing three genera.

| Phylogenomic metrics | Clade I-c versus |  |  |
| --- | --- | --- | --- |
|  | *Parageobacillus* | *Anoxybacillus* | *Geobacillus* |
| Average nucleotide identity (mean %) | 77.40 | 77.90 | 76.54 |
| Average amino acid identity (mean %) | 75.27 | 70.73 | 70.30 |

**Table 2.**
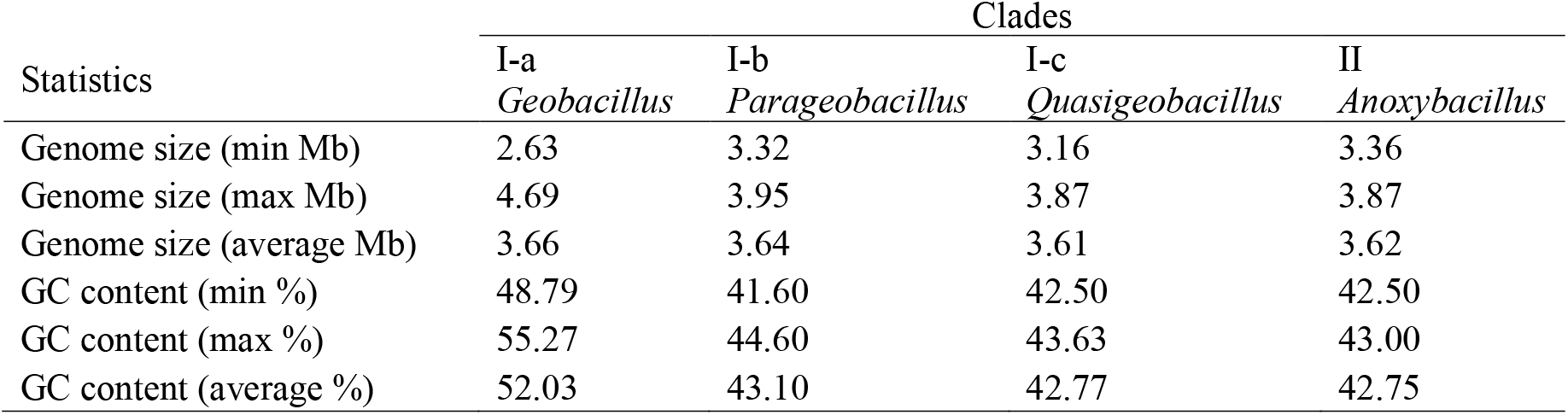
Statistics comparing minimum, maximum and mean values of genome size and GC content per clade.

**Table 3.**
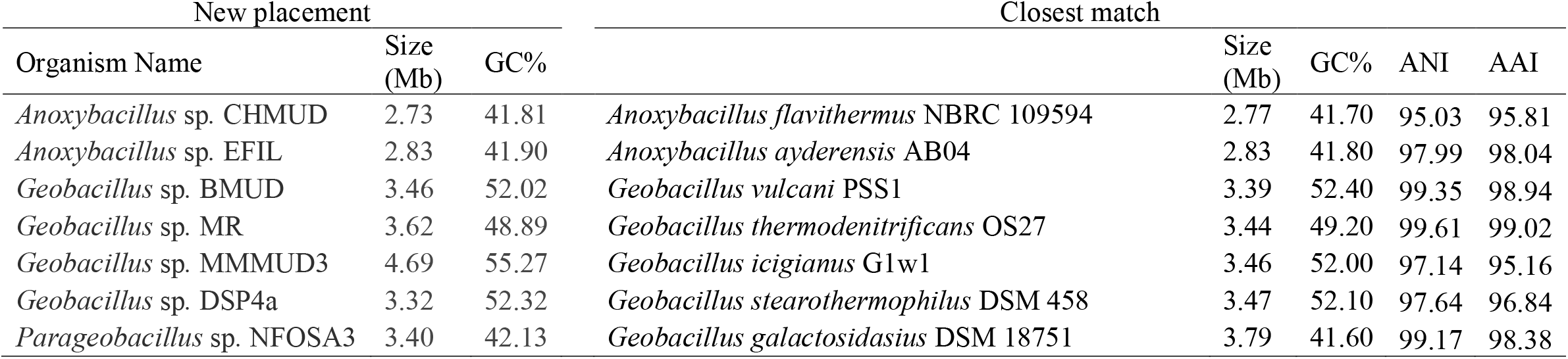
Placement of seven newly sequenced genomes from the study of thermophilic bacteria in Australia.

| New placement |  |  | Closest match |  |  |  |  |
| --- | --- | --- | --- | --- | --- | --- | --- |
| Organism Name | Size (Mb) | GC% |  | Size (Mb) | GC% | ANI | AAI |
| *Anoxybacillus* sp. CHMUD | 2.73 | 41.81 | *Anoxybacillus flavithermus* NBRC 109594 | 2.77 | 41.70 | 95.03 | 95.81 |
| *Anoxybacillus* sp. EFIL | 2.83 | 41.90 | *Anoxybacillus ayderensis* AB04 | 2.83 | 41.80 | 97.99 | 98.04 |
| *Geobacillus* sp. BMUD | 3.46 | 52.02 | *Geobacillus vulcani* PSS1 | 3.39 | 52.40 | 99.35 | 98.94 |
| *Geobacillus* sp. MR | 3.62 | 48.89 | *Geobacillus thermodenitrificans* OS27 | 3.44 | 49.20 | 99.61 | 99.02 |
| *Geobacillus* sp. MMMUD3 | 4.69 | 55.27 | *Geobacillus icigianus* G1w1 | 3.46 | 52.00 | 97.14 | 95.16 |
| *Geobacillus* sp. DSP4a | 3.32 | 52.32 | *Geobacillus stearothermophilus* DSM 458 | 3.47 | 52.10 | 97.64 | 96.84 |
| *Parageobacillus* sp. NFOSA3 | 3.40 | 42.13 | *Geobacillus galactosidasius* DSM 18751 | 3.79 | 41.60 | 99.17 | 98.38 |

### Genes unique to clade I-c

A total of 6 CDS were found to be unique to clade I-c. Names, descriptions and products of CDS are shown in **Table 4**, and DNA sequences for each CDS are shown in **Table 5**.

**Table 4.** Description names of genes unique to clade I-c.

| CDS | Description | Product | Protein ID |
| --- | --- | --- | --- |
| N667_RS0109915 | Protein Homology | MFS transporter | WP_027409424.1 |
| N667_RS0100490 | Protein Homology | spore coat protein | WP_027407730.1 |
| N667_RS0109235 | GeneMarkS-2+ | hypothetical protein | WP_027409309.1 |
| N667_RS0101915 | Protein Homology | MurR/RpiR family transcriptional regulator | WP_051529925.1 |
| N667_RS0106250 | N/A* | hypothetical protein [52] | N/A* |
| N667_RS0109695 | Protein Homology | FUSC family protein | WP_027409385.1 |

\*N/A = Not available

**Table 5.**
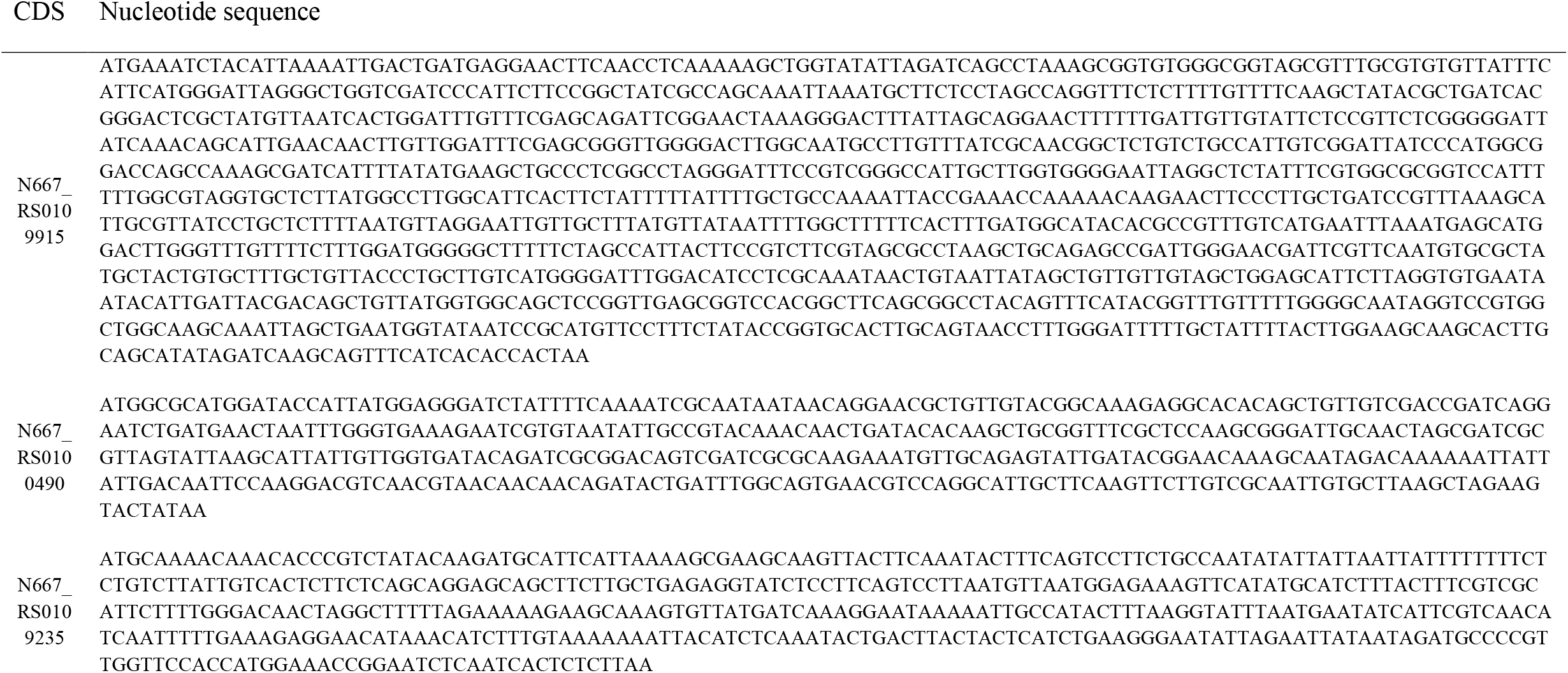
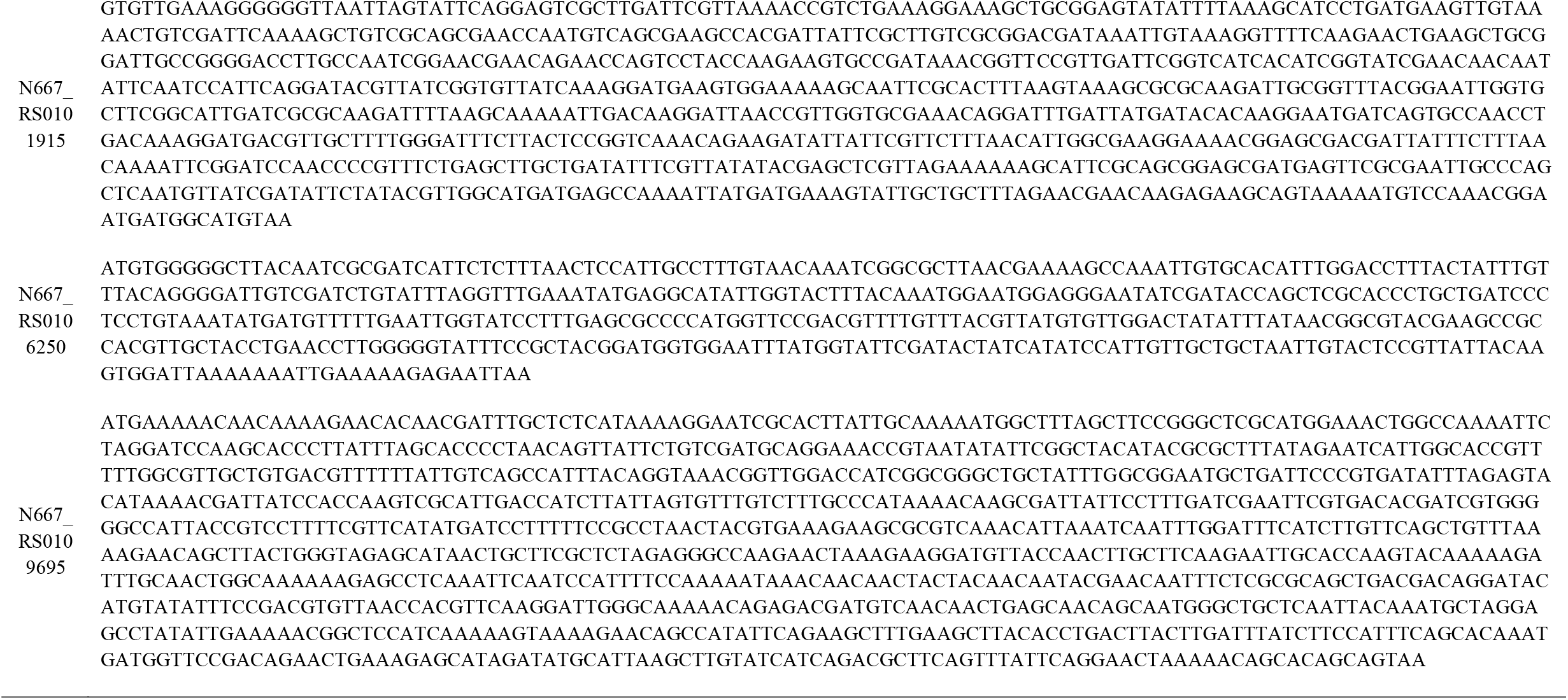
Nucleotide sequences of genes unique to clade I-c.

**Table 6** shows the BLASTn alignment results of these CDS against all members of clade I-c showing that the 6 genes are present in all members of the clade. **Table 7** shows the best hit from the BLASTn results to genomes not present in clade I-c. In all cases, the best hit outside of clade I-c is a genome assembly distant from *Anoxybacillus, Geobacillus* and *Parageobacillus*. The percentage identity and the percentage alignment overlap of the best hit to assemblies outside of clade I-c are presented in the table. For these 6 genes, the closest genomes outside of clade I-c were *Bacillus methanolicus, Bacillus* sp., *Polaribacter* sp., *Paenibacillus* sp., *Bacillus alveayuensis* and *Mesobacillus jeotgali*.

**Table 6.** Results of BLASTn alignment of 6 CDS unique to clade I-c against genomes of clade I-c.

| CDS | subject accession | subject title | % identity | evalue | bitscore | % Overlap* |
| --- | --- | --- | --- | --- | --- | --- |
| N667_RS0109915 | NZ_JHVN01000010.1 | *Anoxybacillus tepidamans* PS2 N667 | 100.0 | 0 | 2255 | 1.00 |
| N667_RS0109915 | NZ_NASY01000037.1 | *Anoxybacillus* sp. UARK-01 Contig37 | 77.3 | 1.97E-177 | 621 | 0.87 |
| N667_RS0109915 | NZ_LPUG01000023.1 | *Anoxybacillus* sp. P3H1B AT864_contig000023 | 77.2 | 9.17E-176 | 616 | 0.87 |
| N667_RS0109915 | NZ_CP015435.1 | *Anoxybacillus* sp. B2M1 | 76.9 | 9.23E-171 | 599 | 0.87 |
| N667_RS0109915 | NZ_CP015436.1 | *Anoxybacillus* sp. B7M1 | 76.9 | 9.23E-171 | 599 | 0.88 |
| N667_RS0109695 | NZ_JHVN01000010.1 | *Anoxybacillus tepidamans* PS2 N667 | 100.0 | 0 | 1951 | 1.00 |
| N667_RS0109695 | NZ_NASY01000037.1 | *Anoxybacillus* sp. UARK-01 Contig37 | 73.9 | 4.12E-94 | 344 | 0.84 |
| N667_RS0109695 | NZ_CP015436.1 | *Anoxybacillus* sp. B7M1 | 73.6 | 4.15E-89 | 327 | 0.84 |
| N667_RS0109695 | NZ_CP015435.1 | *Anoxybacillus* sp. B2M1 | 73.6 | 4.15E-89 | 327 | 0.84 |
| N667_RS0109695 | NZ_LPUG01000023.1 | *Anoxybacillus* sp. P3H1B AT864_contig000023 | 72.7 | 1.15E-89 | 329 | 1.00 |
| N667_RS0109235 | NZ_JHVN01000009.1 | *Anoxybacillus tepidamans* PS2 N667 | 100.0 | 0 | 870 | 1.00 |
| N667_RS0109235 | NZ_CP015435.1 | *Anoxybacillus* sp. B2M1 | 77.8 | 1.57E-05 | 49.1 | 0.17 |
| N667_RS0109235 | NZ_NASY01000046.1 | *Anoxybacillus* sp. UARK-01 Contig46 | 76.2 | 3.37E-07 | 54.7 | 0.22 |
| N667_RS0109235 | NZ_LPUG01000025.1 | *Anoxybacillus* sp. P3H1B AT864_contig000025 | 76.2 | 3.37E-07 | 54.7 | 0.22 |
| N667_RS0109235 | NZ_CP015436.1 | *Anoxybacillus* sp. B7M1 | 75.2 | 1.57E-05 | 49.1 | 0.22 |
| N667_RS0106250 | NZ_JHVN01000004.1 | *Anoxybacillus tepidamans* PS2 N667 | 100.0 | 0 | 854 | 1.00 |
| N667_RS0106250 | NZ_NASY01000026.1 | *Anoxybacillus* sp. UARK-01 Contig26 | 87.2 | 1.99E-04 | 45.4 | 0.08 |
| N667_RS0106250 | NZ_CP015435.1 | *Anoxybacillus* sp. B2M1 | 86.2 | 5.6 | 30.7 | 0.06 |
| N667_RS0106250 | NZ_NASY01000047.1 | *Anoxybacillus* sp. UARK-01 Contig47 | 86.2 | 5.6 | 30.7 | 0.06 |
| N667_RS0106250 | NZ_LPUG01000009.1 | *Anoxybacillus* sp. P3H1B AT864_contig000009 | 86.2 | 5.6 | 30.7 | 0.06 |
| N667_RS0106250 | NZ_LPUG01000010.1 | *Anoxybacillus* sp. P3H1B AT864_contig000010 | 85.7 | 1.99E-04 | 45.4 | 0.09 |
| N667_RS0106250 | NZ_CP015436.1 | *Anoxybacillus* sp. B7M1 | 80.4 | 0.12 | 36.2 | 0.10 |
| N667_RS0101915 | NZ_JHVN01000001.1 | *Anoxybacillus tepidamans* PS2 N667 | 100.0 | 0 | 1596 | 1.00 |
| N667_RS0101915 | NZ_NASY01000051.1 | *Anoxybacillus* sp. UARK-01 Contig51 | 76.8 | 8.87E-130 | 462 | 0.97 |
| N667_RS0101915 | NZ_LPUG01000011.1 | *Anoxybacillus* sp. P3H1B AT864_contig000011 | 76.5 | 1.92E-126 | 451 | 0.97 |
| N667_RS0101915 | NZ_CP015435.1 | *Anoxybacillus* sp. B2M1 | 76.2 | 1.50E-122 | 438 | 0.96 |
| N667_RS0101915 | NZ_CP015436.1 | *Anoxybacillus* sp. B7M1 | 76.2 | 1.50E-122 | 438 | 0.97 |
| N667_RS0100490 | NZ_JHVN01000001.1 | *Anoxybacillus tepidamans* PS2 N667 | 100.0 | 0 | 798 | 1.00 |
| N667_RS0100490 | NZ_NASY01000011.1 | *Anoxybacillus* sp. UARK-01 Contig11 | 89.3 | 0.11 | 36.2 | 0.06 |
| N667_RS0100490 | NZ_CP015436.1 | *Anoxybacillus* sp. B7M1 | 89.3 | 0.11 | 36.2 | 0.06 |
| N667_RS0100490 | NZ_CP015435.1 | *Anoxybacillus* sp. B2M1 | 89.3 | 0.11 | 36.2 | 0.06 |
| N667_RS0100490 | NZ_LPUG01000013.1 | *Anoxybacillus* sp. P3H1B AT864_contig000013 | 89.3 | 0.11 | 36.2 | 0.06 |

\*Calculated dividing length of query by the minimum of either subject length or query length.

**Table 7.**
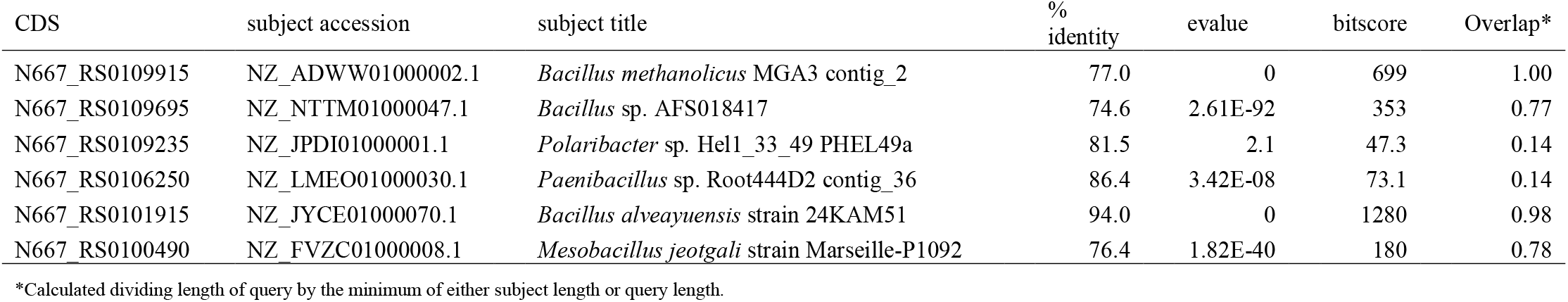
Results of BLASTn alignment of 6 CDS unique to clade I-c against the RefSeq database showing the best hit to genomes outside of clade I-c.

## Discussion

In this paper, we have found that the current taxonomic delimitations of the genera *Anoxybacillus, Geobacillus* and *Parageobacillus* does not represent accurately the phylogenetic relationships within this group.

### Assessment of clade I-c

In the core gene maximum likelihood tree, clade I, which contains the genera *Geobacillus* and *Parageobacillus* in subclades (clade I-a and clade I-b respectively), also contained a third clade (clade I-c) which consisted of genomes ascribed to the genus *Anoxybacillus*. Two further *Anoxybacillus* species present within clade I, *A. vitaminiphilus* and *A. amylolyticus*, exhibited incongruous placement when the core gene tree was compared with the UPGMA dendrograms derived from ANI and AAI similarity matrices of all taxa. The affinities of these two species were difficult to determine from the current evidence and are considered as unresolved taxa. The other *Anoxybacillus* genomes present within clade 1-c, excluding the two species with incongruous placement, were *A. tepidamans, Anoxybacillus* sp. B2M1, *Anoxybacillus* sp. B7M1, *Anoxybacillus* sp. P3H1B and *Anoxybacillus* sp. UARK 01. These genomes were located within clade I-c in each of the three phylogenetic trees, forming a grouping which was consistently distinct from *Geobacillus* (clade I-a) and *Parageobacillus* (clade I-b). Internally, this grouping consisted of two further divisions, with *A. tepidamans* on the one hand, and the remaining four genomes which are closely placed on all of the trees, having ANI values ranging from 98.5 – 99.11% and AAI values ranging from 98.13 – 99.98%. These four genomes can be considered to represent a single genomospecies. The minimum ANI and AAI values from *A. tepidamans* to the members of the genomospecies is 79.48% and 81.72% respectively. The members of clade I-c are therefore considered to consist of two species, *A. tepidamans* and one genomospecies. Due to the consistent separation of the two species in clade I-c from *Geobacillus, Parageobacillus*, and the other members of *Anoxybacillus* in clade II, we consider that the evidence suggests a fourth, new genus is required to accommodate these two species. The average ANI and AAI values separating the candidate new genus from the members of the existing three genera are shown in **Table 1**. The mean ANI and AAI values suggest that the candidate new genus is somewhat more closely related to *Parageobacillus*, followed by *Anoxybacillus* and *Geobacillus* in descending order. The evidence from genome size and GC content also suggests a closer affinity to *Parageobacillus* (**Table 2**). The structure of the three phylogenetic trees reaffirms the distinctiveness of the two species consistently present in clade I-c with respect to the three existing genera. An alternative approach may be to merge these two species with *Parageobacillus*; however, if this course of action was followed, the evidence from the phylogenetic trees would also support the need for *Parageobacillus* to be merged again with *Geobacillus*. Considering the evidence of the core gene maximum likelihood tree, ANI and AAI UPGMA dendrograms and similarity values, we suggest that *A. tepidamans* and the one genomospecies present in clade I-c should be placed into a new genus called *Quasigeobacillus*. Due to the incongruous positioning of *A. vitaminiphilus* and *A. amylolyticus* the affinities of these two species remains unresolved at this point.

### Genes unique to clade I-c

The core gene maximum likelihood tree, ANI and AAI similarity matrices indicate that clade I-c is distinct from clade I-a and clade I-b. To assess the potential presence of distinct genes within clade I-c, a BLASTn alignment of all CDS from clade I-c was performed. CDS which were found to be present in all members of clade I-c and no members of clade I-a, clade I-b and clade II, on the basis of the BLAST alignments were selected. This analysis revealed 6 genes unique to clade I-c. In all cases, when the 6 CDS were aligned using BLASTn against the NCBI RefSeq database, the nearest matching assemblies excluding clade I-c were all from taxa distinct from *Anoxybacillus, Geobacillus* and *Parageobacillus*. The origin of these 6 genes in clade I-c may derive from a horizontal gene transfer to the basal ancestor of clade I-c.

### Assessment of *Parageobacillus*

Clade I-b contains inconsistent naming at the genus level. Six of the genomes that have been assigned to *Geobacillus* and *Anoxybacillus* have been found to belong to *Parageobacillus* in clade I-b. These genomes have not been named to species level or validly described. The list includes one genome named as *A. flavithermus* strain B4168 and five *Geobacillus* genomes, including *Geobacillus* sp. 44B, *Geobacillus* sp. Y41MC1, *Geobacillus* sp. WCH70, *Geobacillus* sp. LYN3 and *Geobacillus* sp. 44C. The genome named *A. flavithermus* strain B4168 is incorrectly assigned at the genus and species level.

## Conclusions

The process of identifying and naming isolates after whole-genome sequencing is usually based on an approach in which the closest genome is identified using multiple methods for alignment of nucleotide or protein sequences and the distance from the nearest genome then determines whether the strain is a new species or belong to an existing species. Although phylogenetic assessment has been recognised as an essential step in this process, the breadth of species assessed together requires further consideration. A combined phylogenomic analysis of the genera *Geobacillus, Parageobacillus* and *Anoxybacillus* has revealed problems in the delimitation of these genera, which would not be revealed if each were considered alone. In the context of determining affinities of a new genome, it is essential to consider closely related species together for construction of a phylogeny. When this process occurs in the absence of a broader phylogenetic assessment, the genus designation for new strains can become incongruent with the phylogeny of the underlying complete set of available genomes. For example, the genomes belonging to clade I-c have been previously assigned to the genus *Anoxybacillus*, and in one case, previously assigned to *Geobacillus*, however, when the two genera are considered together in a phylogenetic analysis, clade I-c is seen to be outside of *Anoxybacillus* and *Geobacillus*.

### Proposed taxonomic changes

#### Description of *Quasigeobacillus* gen. nov

*Quasigeobacillus* (Qua.si.ge.o.ba.cil’lus. L. adv. *quasi* almost like; N.L. masc. n *Geobacillus* a bacterial genus; N.L. masc. n. *Quasigeobacillus*, similar to *Geobacillus*)

The genomes in clade I-c form a distinct group based on genome characteristics and can be separated from *Anoxybacillus, Geobacillus* and *Parageobacillus* on this basis. The position of this clade in relation to the other species is shown in **Figure 2** and **Figure 3**, the ANI and AAI similarity values are shown as a heatmap in **Figure 4** and the full data set for this heatmap is included in the **Supplementary Table S5**. The genome size of this clade ranges from 3.16 to 3.87 Mb and has a GC content ranging from 42.50 to 43.63 %. Therefore, we propose a new genus consisting of *Q. tepidamans*.

**Figure 4.**
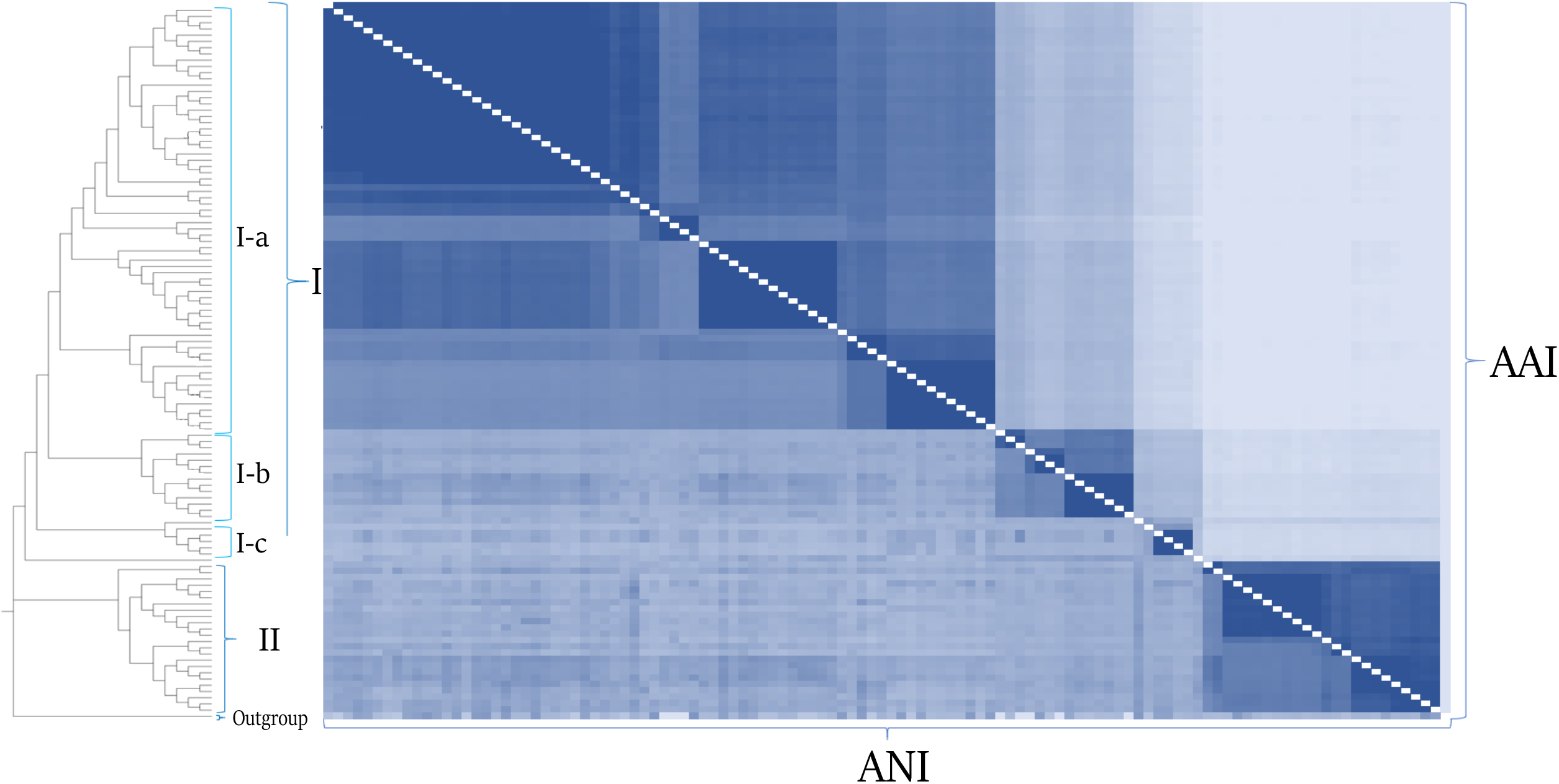

The type species is *Quasigeobacillus tepidamans*.

#### Description of *Quasigeobacillus tepidamans* comb. nov

Etymology *tepidamans*: te.pid.a’mans. L. masc. adj. *tepidus* moderately warm, lukewarm; L. pres. part. *amans* loving; N.L. part. adj. *tepidamans* lowing warm (conditions).

Basonym: *Anoxybacillus tepidamans* Schäffer et al. 2004; *Anoxybacillus tepidamans* Coorevits et al. 2012.

The description for Quasigeobacillus *tepidamans* comb. nov. follows that of *Geobacillus tepidamans* [22], *Anoxybacillus tepidamans* Coorevits et al. [27] and the genus *Quasigeobacillus*.

The type strain is *Quasigeobacillus tepidamans*.

This is available in two culture collections deposits. The American Type Culture Collection ATCC® BAA-942 and Leibniz Institute DSMZ-German Collection of Microorganisms and Cell Cultures GmbH DSM No: 16325, Type Strain, Designation GS5-97, R-35643.

## Supporting information

List_of_Figures

List_of_Tables

Proof of culture collection deposit 1

Proof of culture collection deposit 2

## Availability of data and materials

The data supporting our findings are contained within the manuscript and supplementary materials.

## Author contributions

**Berenice Talamantes-Becerra:** Conceptualization, project administration, Methodology, Data curation, visualization, Formal Analysis, Writing-Original draft preparation and Editing. **Jason Carling:** Conceptualization, Methodology, Visualization, Formal Analysis, Validation, Writing-Original draft preparation and Editing. **Jochen Blom**: Software. **Arthur Georges:** Conceptualization, Resources, Writing-Reviewing and Editing.

## Ethical approval

Not applicable.

## Consent for publication

Not applicable.

## Acknowledgements

BTB would like to acknowledge Josefina Talamantes-Becerra, Angel Medina and Colash.

## Abbreviations

ANI: Average Nucleotide Identity
AAI: Average Amino acid Identity
DDH: DNA-DNA hybridization
CDS: Coding Sequence

