## Supplementary material for "Phylogenetic study of thermophilic genera *Anoxybacillus, Geobacillus, Parageobacillus,* and proposal of a new classification *Quasigeobacillus* gen. nov": List_of_Figures

**List of figures for:**

**List of figures:**

**Figure 1.** Maximum likelihood tree built from alignment of 662 core genes from 113 genomes of *Anoxybacillus*, *Geobacillus*, *Parageobacillus* and one genome of *Bacillus subtilis subsp. spizizenii* TU B 10 selected as an outgroup. Bootstrap values (1000 repetitions) less than 100% are shown on the tree. Scale bar for branch lengths represents 0.1 substitutions per site. Clades and subclades are shown in addition to genome size (Mb) and GC content (%) for all genomes. Two genomes with incongruous positioning in relation to Figure 2 and Figure 3 are shown in green text.

**Figure 2.** UPGMA dendrogram constructed from average nucleotide identity (ANI) similarity matrix for all genomes. Clades equivalent to those in Figure 1 are shown. Scale bar for branch length represents 0.01 substitutions per site. Two genomes with incongruous positioning in relation to Figure 1 and Figure 3 are shown in green text.

**Figure 3.** UPGMA dendrogram constructed from average amino acid identity (AAI) similarity matrix for all genomes. Clades equivalent to those in Figure 1 are shown. Scale bar for branch length represents 0.01 substitutions per site. Two genomes with incongruous positioning in relation to Figure 1 and Figure 2 are shown in green text.

**Figure 4.** Heatmap representation. Average nucleotide identity (ANI) similarity matrix is shown in lower left triangle and average amino acid identity (AAI) similarity matrix is shown upper right triangle. Maximum likelihood tree constructed from alignment of core genes (Figure 1) is shown in alignment with the matrix including clades and subclades. Samples in the matrix are ordered according to the maximum likelihood tree (Figure 1). Colour scale represents values from 57.01% (light blue) to 100% (dark blue). The full data set for this heatmap is included as a Microsoft Excel File in the supplementary material as Supplementary Table S5.
