## Supplementary_information_Tables_S1_S2 for "Phylogenetic study of thermophilic genera *Anoxybacillus, Geobacillus, Parageobacillus,* and proposal of a new classification *Quasigeobacillus* gen. nov"

***Quasigeobacillus* gen. nov.**

Talamantes-Becerra, B., Carling, J, Bloom, J and Georges, A

**Table of Contents:**

| **Supplementary Table S1.** List of selected GenBank genomes for performing the phylogenetic study of thermophilic genera *Anoxybacillus*, *Geobacillus*, *Parageobacillus* and selected genomes of NCBI BioProject PRJNA516322. Genomes are classified by clades according to their position in the maximum likelihood tree (Figure 1) including relevant assembly statistics. | Page 2 |
| --- | --- |
| **Supplementary Table S2.** List of GenBank genome assemblies excluded from the phylogenetic study of thermophilic genera *Anoxybacillus*, *Geobacillus* and *Parageobacillus*. | Page 7 |
| **Supplementary Table S3.** Average nucleotide identity (ANI) similarity matrix for all genomes. Color scale represents values from 66.31% (light blue) to 100% (dark blue). Values are included in all cells of the matrix. | Excel Sheet 1 |
| **Supplementary Table S4.** Average amino acid identity (AAI) similarity matrix for all genomes. Color scale represents values from 57.01% (light blue) to 100% (dark blue). Values are included in all cells of the matrix. | Excel Sheet 2 |
| **Supplementary Table S5.** Average nucleotide identity (ANI) similarity matrix is shown in lower left triangle and average amino acid identity (AAI) similarity matrix is shown upper right triangle. Maximum likelihood tree constructed from alignment of core genes (Figure 1) is shown in alignment with the matrix including clades and subclades. Samples in the matrix are ordered according to the maximum likelihood tree (Figure 1). Color scale represents values from 57.01% (light blue) to 100% (dark blue). Values are included in all cells of the matrix. | Excel Sheet 3 |

**Supplementary Table S1. List of selected NCBI genomes for performing the phylogenetic study of thermophilic genera *Anoxybacillus*, *Geobacillus, Parageobacillus* and selected genomes of NCBI BioProject PRJNA516322. Genomes classified by clades according to their location in the dendrogram, including its relevant statistics and assembly level.**

| **CLADE** | **Organism Name** | **Strain** | **BioSample** | **BioProject** | **Assembly** | **Level** | **Size (Mb)** | **GC%** |
| --- | --- | --- | --- | --- | --- | --- | --- | --- |
| I-a | *Geobacillus genomosp* | JF8 | SAMN02603868 | PRJNA208285 | GCA_000445995.2 | Complete | 3.49 | 52.82 |
| I-a | *Geobacillus icigianus* | G1w1 | SAMN02850065 | PRJNA246135 | GCA_000750005.1 | Contig | 3.46 | 52.00 |
| I-a | *Geobacillus jurassicus* | NBRC 107829 | SAMD00045741 | PRJDB428 | GCA_001544315.1 | Contig | 3.45 | 52.20 |
| I-a | *Geobacillus kaustophilus* | Et2/3 | SAMN03025779 | PRJNA260741 | GCA_000948165.1 | Contig | 3.51 | 51.80 |
| I-a | *Geobacillus kaustophilus* | Et7/4 | SAMN03025780 | PRJNA260742 | GCA_000948285.1 | Contig | 3.68 | 51.70 |
| I-a | *Geobacillus kaustophilus* | GBlys | SAMD00036748 | PRJDB1127 | GCA_000415905.1 | Contig | 3.54 | 52.10 |
| I-a | *Geobacillus kaustophilus* | HTA426 | SAMD00061072 | PRJNA13233 | GCA_000009785.1 | Complete | 3.59 | 51.99 |
| I-a | *Geobacillus kaustophilus* | NBRC 102445 | SAMD00000360 | PRJDB414 | GCA_000739955.1 | Contig | 3.45 | 52.00 |
| I-a | *Geobacillus lituanicus* | N-3 | SAMN05894097 | PRJNA347631 | GCA_002243605.1 | Complete | 3.50 | 52.17 |
| I-a | *Geobacillus proteiniphilus* | 1017 | SAMN06043560 | PRJNA353982 | GCA_001908025.1 | Contig | 3.57 | 51.80 |
| I-a | *Geobacillus* sp. | 46C-IIa | SAMN06347030 | PRJNA354604 | GCA_002077765.1 | Scaffold | 3.47 | 52.10 |
| I-a | *Geobacillus* sp*.* | 47C-IIb | SAMN06347031 | PRJNA354604 | GCA_002077775.1 | Scaffold | 3.35 | 49.60 |
| I-a | *Geobacillus* sp*.* | FJAT-46040 | SAMN07212131 | PRJNA390033 | GCA_002335725.1 | Scaffold | 3.36 | 52.30 |
| I-a | *Geobacillus* sp*.* | B4113_201601 | SAMN04390296 | PRJNA270597 | GCA_001587475.1 | Scaffold | 3.65 | 51.30 |
| I-a | *Geobacillus* sp*.* | C56-T3 | SAMN02598534 | PRJNA41701 | GCA_000092445.1 | Complete | 3.65 | 52.50 |
| I-a | *Geobacillus* sp*.* | CAMR5420 | SAMN02715734 | PRJNA243324 | GCA_000691465.1 | Contig | 3.50 | 51.90 |
| I-a | *Geobacillus* sp*.* | FW23 | SAMN02687994 | PRJNA241053 | GCA_000617945.1 | Contig | 3.49 | 52.20 |
| I-a | *Geobacillus* sp*.* | GHH01 | SAMN02603276 | PRJNA180997 | GCA_000336445.1 | Complete | 3.58 | 52.30 |
| I-a | *Geobacillus* sp*.* | LEMMY01 | SAMN06309877 | PRJNA373874 | GCA_002042905.1 | Contig | 3.59 | 51.90 |
| I-a | *Geobacillus* sp*.* | Manikaran-105 | SAMN08056006 | PRJNA354604 | GCA_002809955.1 | Scaffold | 3.19 | 52.50 |
| I-a | *Geobacillus* sp*.* | MAS1 | SAMN02371536 | PRJNA222590 | GCA_000498995.1 | Scaffold | 3.50 | 52.20 |
| I-a | *Geobacillus* sp*.* | PA-3 | SAMN03963308 | PRJNA292061 | GCA_001412125.1 | Contig | 3.65 | 48.90 |
| I-a | *Geobacillus* sp*.* | Sah69 | SAMN04161460 | PRJNA298658 | GCA_001414205.1 | Contig | 2.99 | 52.60 |
| I-a | *Geobacillus* sp*.* | T6 | SAMN03734430 | PRJNA284908 | GCA_001025095.1 | Contig | 3.66 | 51.90 |
| I-a | *Geobacillus* sp*.* | WSUCF-018B | SAMN08056005 | PRJNA354604 | GCA_002809985.1 | Scaffold | 3.23 | 52.50 |
| I-a | *Geobacillus* sp*.* | Y412MC52 | SAMN00713574 | PRJNA30797 | GCA_000174795.2 | Complete | 3.67 | 52.31 |
| I-a | *Geobacillus* sp*.* | Y412MC61 | SAMN00017500 | PRJNA30537 | GCA_000024705.1 | Complete | 3.67 | 52.31 |
| I-a | *Geobacillus* sp*.* | ZGt-1 | SAMN03420899 | PRJNA278519 | GCA_001026865.1 | Scaffold | 3.48 | 52.10 |
| I-a | *Geobacillus* sp*.* | BMUD | SAMN10787408 | PRJNA516322 | SDLA00000000 | Contig | 3.46 | 52.02 |
| I-a | *Geobacillus* sp*.* | MR | SAMN10787385 | PRJNA516322 | SDLB00000000 | Contig | 3.62 | 48.89 |
| I-a | *Geobacillus* sp*.* | MMMUD3 | SAMN10787384 | PRJNA516322 | SDLC00000000 | Contig | 4.69 | 55.27 |
| I-a | *Geobacillus* sp*.* | DSP4a | SAMN10787358 | PRJNA516322 | SDLD00000000 | Contig | 3.32 | 52.32 |
| I-a | *Geobacillus stearothermophilus* | A1 | SAMN03651168 | PRJNA282772 | GCA_001183895.1 | Scaffold | 3.01 | 52.00 |
| I-a | *Geobacillus stearothermophilus* | B4109 | SAMN03267293 | PRJNA270597 | GCA_001587495.1 | Scaffold | 2.78 | 52.50 |
| I-a | *Geobacillus stearothermophilus* | B4114 | SAMN03267294 | PRJNA270597 | GCA_001587395.1 | Scaffold | 2.77 | 52.80 |
| I-a | *Geobacillus stearothermophilus* | D1 | SAMN03651170 | PRJNA282772 | GCA_001183885.1 | Scaffold | 2.97 | 52.20 |
| I-a | *Geobacillus stearothermophilus* | DSM 458 | SAMN05300636 | PRJNA327158 | GCA_002300135.1 | Complete | 3.47 | 52.10 |
| I-a | *Geobacillus stearothermophilus* | GS27 | SAMN04532070 | PRJNA314192 | GCA_001651555.1 | Scaffold | 2.70 | 52.50 |
| I-a | *Geobacillus stearothermophilus* | P3 | SAMN03651169 | PRJNA282772 | GCA_001183915.1 | Scaffold | 3.02 | 52.00 |
| I-a | *Geobacillus stearothermophilus* | 10 | SAMN04012428 | PRJNA252389 | GCA_001274575.1 | Complete | 3.68 | 52.61 |
| I-a | *Geobacillus stearothermophilus* | ATCC 12980 | SAMN03291237 | PRJNA212538 | GCA_001277805.1 | Scaffold | 2.63 | 53.10 |
| I-a | *Geobacillus stearothermophilus* | ATCC 7953 | SAMN02569809 | PRJNA233505 | GCA_000705495.1 | Contig | 2.79 | 52.40 |
| I-a | *Geobacillus subterraneus* | K | SAMN04633529 | PRJNA318163 | GCA_001632595.1 | Contig | 3.30 | 52.30 |
| I-a | *Geobacillus subterraneus* | KCTC 3922 | SAMN04445793 | PRJNA310054 | GCA_001618685.1 | Complete | 3.47 | 52.20 |
| I-a | *Geobacillus subterraneus* | PSS2 | SAMN02744825 | PRJNA214283 | GCA_000744755.1 | Contig | 3.75 | 51.60 |
| I-a | *Geobacillus thermocatenulatus* | BGSC 93A1 | SAMN06768585 | PRJNA383645 | GCA_002217655.1 | Contig | 3.56 | 51.80 |
| I-a | *Geobacillus thermocatenulatus* | KCTC 3921 | SAMN06016323 | PRJNA353561 | GCA_002243665.1 | Complete | 3.74 | 51.90 |
| I-a | *Geobacillus thermocatenulatus* | GS-1 | SAMN02592613 | PRJNA233553 | GCA_000612265.1 | Contig | 3.52 | 52.10 |
| I-a | *Geobacillus thermodenitrificans* | G11MC16 | SAMN02441802 | PRJNA28341 | GCA_000173035.1 | Contig | 3.55 | 48.80 |
| I-a | *Geobacillus thermodenitrificans* | JSC_T9a | SAMN08056007 | PRJNA354604 | GCA_002810005.1 | Scaffold | 3.51 | 48.90 |
| I-a | *Geobacillus thermodenitrificans* | KCTC3902 | SAMN05894115 | PRJNA347632 | GCA_002072065.1 | Complete | 3.51 | 49.10 |
| I-a | *Geobacillus thermodenitrificans* | OS27 | SAMN08135953 | PRJNA421250 | GCA_003061505.1 | Contig | 3.44 | 49.20 |
| I-a | *Geobacillus thermodenitrificans* | T12 | SAMN06459234 | PRJNA377291 | GCA_002119625.1 | Complete | 3.76 | 48.79 |
| I-a | *Geobacillus thermodenitrificans* | NG80-2 | SAMN02603899 | PRJNA18655 | GCA_000015745.1 | Complete | 3.61 | 48.85 |
| I-a | *Geobacillus thermodenitrificans* subsp*. thermodenitrificans* | DSM 465 | SAMN02386948 | PRJNA224955 | GCA_000496575.1 | Contig | 3.40 | 49.00 |
| I-a | *Geobacillus thermoleovorans* | FJAT-2391 | SAMN04631815 | PRJNA340206 | GCA_001719205.1 | Complete | 3.54 | 52.20 |
| I-a | *Geobacillus thermoleovorans* | ID-1 | SAMN05893589 | PRJNA347629 | GCA_002706565.1 | Complete | 3.77 | 48.98 |
| I-a | *Geobacillus thermoleovorans* | KCTC 3570 | SAMN04455789 | PRJNA310809 | GCA_001610955.1 | Complete | 3.50 | 52.29 |
| I-a | *Geobacillus thermoleovorans* | N7 | SAMN04633480 | PRJNA318151 | GCA_001707765.1 | Contig | 3.40 | 52.40 |
| I-a | *Geobacillus thermoleovorans* | SGAir0734 | SAMN08222721 | PRJNA388547 | GCA_003182675.2 | Complete | 3.69 | 51.96 |
| I-a | *Geobacillus thermoleovorans* | SURF-48B | SAMN06347032 | PRJNA354604 | GCA_002077815.1 | Scaffold | 3.39 | 52.60 |
| I-a | *Geobacillus thermoleovorans* | B23 | SAMD00036769 | PRJDB1475 | GCA_000474195.1 | Contig | 3.35 | 52.30 |
| I-a | *Geobacillus thermoleovorans* | CCB_US3_UF5 | SAMN02603099 | PRJNA76489 | GCA_000236605.1 | Complete | 3.60 | 52.30 |
| I-a | *Geobacillus uzenensis* | BGSC 92A1 | SAMN06768593 | PRJNA383648 | GCA_002217665.1 | Contig | 3.36 | 52.20 |
| I-a | *Geobacillus vulcani* | PSS1 | SAMN02744877 | PRJNA214297 | GCA_000733845.1 | Contig | 3.39 | 52.40 |
| I-a | *Geobacillus zalihae* | SURF-114 | SAMN06347033 | PRJNA354604 | GCA_002077855.1 | Contig | 3.43 | 52.40 |
| I-a | *Geobacillus zalihae* | SURF-189 | SAMN06347034 | PRJNA354604 | GCA_002077835.1 | Scaffold | 3.55 | 52.40 |
| I-a | *Geobacillus zalihae* | NBRC 101842 | SAMD00045737 | PRJDB415 | GCA_001544135.1 | Contig | 3.54 | 51.90 |
| I-b | *Anoxybacillus flavithermus* | B4168 | SAMN03267297 | PRJNA270597 | GCA_001587555.1 | Scaffold | 3.72 | 43.80 |
| I-b | *Geobacillus* sp*.* | 44B | SAMN06347028 | PRJNA354604 | GCA_002077755.1 | Contig | 3.54 | 44.60 |
| I-b | *Geobacillus* sp*.* | 44C | SAMN06347029 | PRJNA354604 | GCA_002077865.1 | Scaffold | 3.22 | 42.30 |
| I-b | *Geobacillus* sp*.* | LYN3 | SAMN08933550 | PRJNA450255 | GCA_003064505.1 | Scaffold | 3.24 | 42.60 |
| I-b | *Geobacillus* sp*.* | WCH70 | SAMN00000635 | PRJNA20805 | GCA_000023385.1 | Complete | 3.51 | 42.76 |
| I-b | *Geobacillus* sp*.* | Y4.1MC1 | SAMN00002562 | PRJNA33183 | GCA_000166075.1 | Complete | 3.91 | 44.00 |
| I-b | *Parageobacillus caldoxylosilyticus* | B4119 | SAMN03267290 | PRJNA270597 | GCA_001587505.1 | Scaffold | 3.95 | 44.00 |
| I-b | *Parageobacillus galactosidasius* | DSM 18751 | SAMN06770004 | PRJNA383662 | GCA_002217735.1 | Contig | 3.79 | 41.60 |
| I-b | *Parageobacillus genomosp.* 1 | NUB3621 | SAMN02727286 | PRJNA189971 | GCA_000632515.1 | Complete | 3.62 | 44.40 |
| I-b | *Parageobacillus* sp*.* | NFOSA3 | SAMN10787357 | PRJNA516322 | SDLE00000000 | Contig | 3.40 | 42.13 |
| I-b | *Parageobacillus thermantarcticus* | M1 | SAMN05192569 | PRJEB17059 | GCA_900111865.1 | Scaffold | 3.45 | 43.70 |
| I-b | *Parageobacillus thermoglucosidasius* | DSM 2542 | SAMN04099008 | PRJNA296418 | GCA_001295365.1 | Complete | 3.87 | 43.90 |
| I-b | *Parageobacillus toebii* NBRC 107807 | DSM 14590 | SAMN09062732 | PRJNA455457 | GCA_003688615.1 | Contig | 3.32 | 42.40 |
| I-b | *Parageobacillus yumthangensis* | AYN2 | SAMN07653191 | PRJNA407404 | GCA_002494375.1 | Scaffold | 3.41 | 42.40 |
| I-c | *Anoxybacillus* sp. | b2m1 | SAMN04528779 | PRJNA314078 | GCA_001634265.1 | Complete | 3.82 | 42.50 |
| I-c | *Anoxybacillus* sp. | b7m1 | SAMN04528789 | PRJNA314079 | GCA_001634305.1 | Complete | 3.87 | 42.51 |
| I-c | *Anoxybacillus* sp. | P3H1B | SAMN04324283 | PRJNA305084 | GCA_001560855.1 | Contig | 3.54 | 42.60 |
| I-c | *Anoxybacillus* sp. | UARK-01 | SAMN06624670 | PRJNA379989 | GCA_002075365.1 | Contig | 3.67 | 42.60 |
| I-c | *Anoxybacillus tepidamans* | PS2 | SAMN02952993 | PRJNA214279 | GCA_000620165.1 | Scaffold | 3.36 | 43.00 |
| II | *Anoxybacillus ayderensis* | AB04 | SAMN03003524 | PRJNA258494 | GCA_000833605.1 | Contig | 2.83 | 41.80 |
| II | *Anoxybacillus ayderensis* | MT-Cab | SAMN06675368 | PRJNA236114 | GCA_002117565.1 | Contig | 2.58 | 41.90 |
| II | *Anoxybacillus ayderensis* | SK3-4 | SAMN02471430 | PRJNA174378 | GCA_000443775.1 | Contig | 2.67 | 41.90 |
| II | *Anoxybacillus flavithermus* | 52-1A | SAMN06925341 | PRJNA386099 | GCA_002197485.1 | Complete | 2.83 | 42.21 |
| II | *Anoxybacillus flavithermus* | AF14 | SAMN04530316 | PRJNA314192 | GCA_001651525.1 | Scaffold | 2.56 | 41.80 |
| II | *Anoxybacillus flavithermus* | AF16 | SAMN04532071 | PRJNA314192 | GCA_001651545.1 | Scaffold | 2.65 | 41.10 |
| II | *Anoxybacillus flavithermus* | DSM 2641T | SAMN06758772 | PRJNA383058 | GCA_002243705.1 | Complete | 2.81 | 41.80 |
| II | *Anoxybacillus flavithermus* | KU2-6_11 | SAMN07831762 | PRJNA415688 | GCA_002742685.1 | Contig | 2.65 | 41.50 |
| II | *Anoxybacillus flavithermus* | WK1 | SAMN02604185 | PRJNA28245 | GCA_000019045.1 | Complete | 2.85 | 41.80 |
| II | *Anoxybacillus flavithermus* | AK1 | SAMN02470127 | PRJNA190633 | GCA_000353425.1 | Contig | 2.63 | 42.70 |
| II | *Anoxybacillus flavithermus* | NBRC 109594 | SAMD00036730 | PRJDB1085 | GCA_000367505.1 | Contig | 2.77 | 41.70 |
| II | *Anoxybacillus flavithermus* | TNO-09.006 | SAMN02471214 | PRJNA169174 | GCA_000327465.1 | Scaffold | 2.66 | 41.80 |
| II | *Anoxybacillus flavithermus* subsp*. yunnanensis* | E13 | SAMN02298030 | PRJNA213809 | GCA_000753835.1 | Scaffold | 2.84 | 41.60 |
| II | *Anoxybacillus gonensis* | DT3-1 | SAMN02471428 | PRJNA182115 | GCA_000346275.1 | Contig | 2.60 | 41.50 |
| II | *Anoxybacillus gonensis* | G2 | SAMN03121471 | PRJNA290998 | GCA_001187595.1 | Complete | 2.80 | 41.70 |
| II | *Anoxybacillus gonensis* | G2 | SAMN03121471 | PRJNA264351 | GCA_000770375.1 | Complete | 2.76 | 41.50 |
| II | *Anoxybacillus kamchatkensis* | G10 | SAMN02470192 | PRJNA170961 | GCA_000283415.1 | Contig | 2.86 | 41.30 |
| II | *Anoxybacillus mongoliensis* | MB4 | SAMN05449942 | PRJNA356103 | GCA_001914435.1 | Contig | 2.81 | 41.70 |
| II | *Anoxybacillus pushchinoensis* | K1 | SAMN05216169 | PRJEB17036 | GCA_900111795.1 | Scaffold | 2.63 | 42.10 |
| II | *Anoxybacillus* sp*.* | CHMUD | SAMN10786853 | PRJNA516322 | SDLG00000000 | Contig | 2.73 | 41.81 |
| II | *Anoxybacillus* sp*.* | EFIL | SAMN10786848 | PRJNA516322 | SDLH00000000 | Contig | 2.83 | 41.90 |
| II | *Anoxybacillus* sp*.* | 103 | SAMN06020137 | PRJNA353767 | GCA_001996285.1 | Scaffold | 2.72 | 41.60 |
| II | *Anoxybacillus suryakundensis* | DSM 27374 | SAMN03840601 | PRJNA288979 | GCA_001418025.1 | Scaffold | 2.59 | 41.50 |
| II | *Anoxybacillus thermarum* | AF/04 | SAMN03078685 | PRJNA260786 | GCA_000836725.1 | Contig | 2.74 | 42.00 |
| Outgroup | *Bacillus subtilis* subsp*. spizizenii* | TU-B-10 | SAMN02603352 | PRJNA68561 | GCA_000227465.1 | Complete | 4.21 | 43.80 |
| (?) | *Anoxybacillus amylolyticus* | DSM 15939 | SAMN04570556 | PRJNA315823 | GCA_001634285.1 | Complete | 3.16 | 43.63 |
| I (?) | *Anoxybacillus vitaminiphilus* | CGMCC 1.8979 | SAMN06296536 | PRJNA370109 | GCA_003259935.1 | Scaffold | 3.56 | 40.10 |

**Supplementary Table S2. List of NCBI genome assemblies excluded from the phylogenetic study of** **thermophilic genera *Anoxybacillus*, *Geobacillus* and *Parageobacillus.***

| **Organism Name** | **Description** | **BioSample** | **BioProject** | **Assembly** |
| --- | --- | --- | --- | --- |
| *Anoxybacillus flavithermus* 25 | Reduced the number of core genes. Excluded from refseq: many frameshifted proteins | SAMN02988316 | PRJNA258119 | GCA_000753775.1 |
| *Anoxybacillus* sp*.* BCO1 | Excluded due to comparatively small number of CDS in the assembly annotations and reduced the number of core genes. | SAMN03075636 | PRJNA261743 | GCA_000787435.1 |
| *Anoxybacillus* sp*.* KU2-6(11) | Excluded due to comparatively small number of CDS in the assembly annotations and reduced the number of core genes. | SAMN02991088 | PRJNA258246 | GCA_000753875.1 |
| *Geobacillus* sp*.* 12AMOR1 | Anomalous assembly. Excluded from refseq: Kimeric | SAMN03400067 | PRJNA277925 | GCA_001028085.1 |
| *Geobacillus* sp*.* 15 | Anomalous assembly. Excluded from refseq: contaminated | SAMN04566681 | PRJNA315614 | GCA_001624615.1 |
| *Geobacillus* sp. 8 | Reduced the number of core genes. It is not *Geobacillus* or *Anoxybacillus.* | SAMN04566680 | PRJNA315613 | GCA_001624605.1 |
| *Geobacillus* sp*.* A8 | Excluded due to comparatively small number of CDS in the assembly annotations and reduced the number of core genes. | SAMN03382575 | PRJNA276936 | GCA_003263855.1 |
| *Geobacillus* sp. BCO2 | Reduced the number of core genes. Excluded from refseq: many frameshift proteins | SAMN03075634 | PRJNA261745 | GCA_001294475.1 |
| *Geobacillus* sp*.* CAMR12739 | Reduced the number of core genes. Excluded from refseq: many frameshift proteins | SAMN02715733 | PRJNA243323 | GCA_000691445.1 |
| *Geobacillus* sp*.* JS12 | Excluded from refseq: many frameshift proteins | SAMN04544377 | PRJNA314835 | GCA_001592395.1 |
| *Geobacillus* sp*.* LC300 | Anomalous assembly. Excluded from refseq: Kimeric | SAMN02903857 | PRJNA254417 | GCA_001191625.1 |
| *Geobacillus* sp*.* WSUCF1 | Reduced the number of core genes. Excluded from refseq: many frameshift proteins | SAMN02471005 | PRJNA192273 | GCA_000422025.1 |
| *Geobacillus stearothermophilus* 22 | Anomalous assembly. Excluded from refseq: contaminated | SAMN02769493 | PRJNA246673 | GCA_000743495.1 |
| *Geobacillus stearothermophilus* 53 | Anomalous assembly. Excluded from refseq: contaminated | SAMN02850071 | PRJNA252422 | GCA_000749985.1 |
| *Geobacillus stearothermophilus* C1BS50MT1 | Anomalous assembly. Excluded from refseq: contaminated | SAMN04378115 | PRJNA305084 | GCA_001620045.1 |
| *Geobacillus* sp*.* ZCTH4_G | Excluded from refseq: derived from metagenome | SAMN08628109 | PRJNA436557 | GCA_003388745.1 |
| *Anoxybacillus suryakundensis* DSM 27374 | Duplicate assembly of GCA_001418025.1 | SAMN03840601 | PRJEB10551 | GCA_001517225.1 |
| *Anoxybacillus geothermalis* ATCC BAA 2555 | It has a genome size two times the average of *Anoxybacillus* and *Geobacillus*. | SAMN03025781 | PRJNA260743 | GCA_001587555.1 |
