## Supplementary material for "Phylogenetic study of thermophilic genera *Anoxybacillus, Geobacillus, Parageobacillus,* and proposal of a new classification *Quasigeobacillus* gen. nov": Proof of culture collection deposit 1

ATCC

Product Sheet

### *Geobacillus tepidamans* (ATCC® BAA-942™)

#### Please read this FIRST

Biosafety Level
1

#### Intended Use

This product is intended for research use only. It is not intended for any animal or human therapeutic or diagnostic use.

#### Citation of Strain

If use of this culture results in a scientific publication, it should be cited in that manuscript in the following manner: *Geobacillus tepidamans* (ATCC® BAA-942™)

#### Description

**Designation:** GS5-97 [DSM 16325]

#### Propagation

**Medium**
ATCC® Medium 18: Trypticase Soy Agar/Broth

**Growth Conditions**
**Temperature:** 55.0°C
**Atmosphere:** Aerobic

#### References

References and other information relating to this product are available online at www.atcc.org.

#### Biosafety Level: 1

Appropriate safety procedures should always be used with this material. Laboratory safety is discussed in the current publication of the *Biosafety in Microbiological and Biomedical Laboratories* from the U.S. Department of Health and Human Services Centers for Disease Control and Prevention and National Institutes for Health.

#### ATCC Warranty

ATCC® products are warranted for 30 days from the date of shipment, and this warranty is valid only if the product is stored and handled according to the information included on this product information sheet. If the ATCC® product is a living cell or microorganism, ATCC lists the media formulation that has been found to be effective for this product. While other, unspecified media may also produce satisfactory results, a change in media or the absence of an additive from the ATCC recommended media may affect recovery, growth and/or function of this product. If an alternative medium formulation is used, the ATCC warranty for viability is no longer valid.

#### Disclaimers

This product is intended for laboratory research purposes only. It is not intended for use in humans.
While ATCC uses reasonable efforts to include accurate and up-to-date information on this product sheet, ATCC makes no warranties or representations as to its accuracy. Citations from scientific literature and patents are provided for informational purposes only. ATCC does not warrant that such information has been confirmed to be accurate.
This product is sent with the condition that you are responsible for its safe storage, handling, and use. ATCC is not liable for any damages or injuries arising from receipt and/or use of this product. While reasonable effort is made to insure authenticity and reliability of materials on deposit, ATCC is not liable for damages arising from the misidentification or misrepresentation of such materials.
Please see the enclosed Material Transfer Agreement (MTA) for further details regarding the use of this product. The MTA is also available on our Web site at www.atcc.org

Additional information on this culture is available on the ATCC web site at www.atcc.org.
© ATCC 2021. All rights reserved. ATCC is a registered trademark of the American Type Culture Collection. [01/01]

American Type Culture Collection
PO Box 1549
Manassas, VA 20108 USA
www.atcc.org

800.638.6597 or 703.365.2700


*Or contact your local distributor*
