## Supplementary material for "Phylogenetic study of thermophilic genera *Anoxybacillus, Geobacillus, Parageobacillus,* and proposal of a new classification *Quasigeobacillus* gen. nov": Proof of culture collection deposit 2

Leibniz Institute DSMZ-German Collection of Microorganisms and Cell Cultures GmbH

### Anoxybacillus tepidamans

DSM 16325

BACTERIA

ⓘ How to read the following data (Example)

**Name:**
***Anoxybacillus tepidamans*** (Schäffer et al. 2004) Coorevits et al. 2012

Synonym(s):
*Geobacillus tepidamans* Schäffer et al. 2004

**DSM No.:**
**16325, Type strain**

Strain designation:
GS5-97, R-35643

Other collection no. or WDCM no.:
ATCC BAA-942

Isolated from:
extraction juice samples

Country:
Austria
Beet Sugar Factory, Leopoldsdorf, lower Austria

Date of sampling:
before 01.04.2004

**Nagoya Protocol Restrictions:**
There are NO known Nagoya Protocol restrictions for this strain.

History:
<- P. Messner; GS5-97 <- M. Graninger and A. Scheberl <- F. Hollaus

Genbank accession numbers:
16S rRNA gene: AY563003
16S rRNA gene: FN428691

Cultivation conditions:
Medium 220 +$MnSO_4$, 55°C

Complete DSMZ Media List

Summary and additional information:
<- P. Messner; GS5-97 <- M. Graninger and A. Scheberl <- F. Hollaus. Extraction juice samples; Austria. **Type strain.** Taxonomy/description (9313, 16936). Sequence accession no. 16S rRNA gene: AY563003 and FN428691. (Medium 220 +$MnSO_4$, 55°C).

For detailed information:
BacDive - The Bacterial Diversity Metadatabase

Literature:
9313, 16936

Risk group:
1 (classification according to German TRBA)

Supplied as:

| Delivery form | Prices |
|---|---|
| Freeze Dried | 90,- € |
| Active culture on request | 210,- € |
| DNA | 135,- € |

Price Category for this culture: **1**

Freight and handling charges will be added. See price list.

Other cultures:
All DSMZ cultures of the species

Print data sheet

Add to Cart

Open Pricelist

#### Help Topics

FAQ →

Order & Delivery →

Safety →

Quality assurance →
